# Discovery of metallophore diversity in *Microbulbifer* in mixed culture with a coral pathogen using computational mass spectrometry and genome mining

**DOI:** 10.64898/2026.03.11.711190

**Authors:** Mónica Monge-Loría, Colin Brady, Hongwei Wu, Allegra Aron, Neha Garg

## Abstract

Iron is an essential component of cellular biology. Thus, iron’s low bioavailability is a key evolutionary pressure guiding microbial dynamics in the marine environment. Among marine bacteria, *Microbulbifer* is an underexplored and functionally versatile bacterial genus, which is commonly associated with sponges, algae, corals and sediments. Previously, genome analyses have revealed that *Microbulbifer* spp. can degrade polymers and synthesize natural products. Despite their recognized potential to produce secondary metabolites, siderophores are yet to be identified in *Microbulbifer*, and their iron acquisition strategies remain largely unknown. Here, we developed a comprehensive mass spectrometry-based query language (MassQL) code to determine siderophore production by *Microbulbifer* spp. in mono- and mixed culture with a marine pathogen, which can be replicated for discovery of these compounds in any organism. Using this workflow, we discovered a new metallophore, which we named bulbichelin, as well as a suite of previously unreported petrobactins containing unprecedented longer chain length acylation on the central spermidine moiety. We applied genome mining methods to describe the biosynthesis of these compounds. Using metal infusion mass spectrometry, we show that bulbichelins bind a variety of metals. Notably, neither of these compounds were produced in a co-culture of *Microbulbifer* with coral-derived pathogen *Vibrio coralliilyticus* Cn52-H1. This observation suggests that *Microbulbifer* uses alternate strategies in a mixed community, such as siderophore piracy for metal acquisition. Understanding how siderophores shape interspecies interactions between *Microbulbifer* spp. and other marine organisms will aid in unraveling the chemical and catalytic versatility of this genus and adaptation in nutrient deplete marine environment.

## Introduction

*Microbulbifer* spp. bacteria, common in marine microbiomes, are recognized for their ability to enzymatically degrade biopolymers such as cellulose, xylan, chitin, agar and alginate^1–7^. Additionally, multiple biosynthetic gene clusters (BGCs) that together encode pathways to produce specialized metabolites have been identified in *Microbulbifer* spp. genomes ^8, 9^. These analyses have revealed the genetic potential of *Microbulbifer* to produce non-ribosomal peptides (NRPs), polyketides, ribosomally synthesized and post-translationally modified peptides (RiPPs), ectoine, butyrolactones, polybrominated compounds, and metallophores^8, 10^. However, only a few compounds have been isolated or identified in this genus, with the subset of *Microbulbifer* secondary metabolites consisting mainly of bulbiferates^11^, bulbiferamides^12, 13^, pseudobulbiferamides^14^ and the chalkophore bulbicupramide^10^. Compounds, such as Bulbiferates A and B (antibacterial acetamidohydroxybenzoates)^11^, are postulated to play key roles in chemical defense and microbial competition^12, 13^, illustrating their function in ecological interactions. Thus, the disparity between the number of BGCs present in *Microbulbifer* genomes and the number of identified compounds encoded by these BGCs highlights a significant gap between genomic potential and identified natural products. This discrepancy presents an opportunity to explore the ecological roles of these compounds in marine microbiomes and discover novel bioactive molecules^9^.

As *Microbulbifer* spp. bacteria are associated with commensal microbiomes, we previously explored production of antimicrobial compounds by *Microbulbifer* against a marine pathogen, *Vibrio coralliilyticus*, isolated from a similar environment^15^. This work led to the serendipitous discovery of the ability of *Microbulbifer* sp. bacteria to enzymatically degrade amphibactin, an iron-acquiring siderophore produced by the pathogen *V. coralliilyticus*. This mechanism, identified via coculture experiments, involves proteolytic cleavage of the peptide backbone, which reduces the siderophore’s iron-binding affinity. This finding suggests that *Microbulbifer* can disrupt the iron acquisition of *V. coralliilyticus*, impacting its survival and proliferation in the host. However, iron acquisition mechanism of *Microbulbifer* itself remains unknown. Iron is an essential element for almost all living organisms, as it is involved in oxygen transport and storage^16^, DNA synthesis and repair^17^, cellular respiration^18^, and electron transport^19, 20^. In addition to its redox properties, the prevalence of iron as a cofactor reflects iron’s abundance and availability during early life development^21, 22^. Therefore, iron influences microbial ecology by acting as a limiting nutrient, a cofactor for essential enzymes, and a driver of electron transport. Competition for iron therefore became an evolutionary driver for the development of iron acquisition strategies especially in aquatic environments where bioavailable iron concentration is low, as introduced below.

The iron concentration in the ocean is at sub nanomolar level whereas its biological requirement spans the nanomolar to micromolar range^23, 24^. This discrepancy prompted the “iron hypothesis”, where iron is proposed to be the limiting nutrient for primary producers in the marine environment^25^. Iron limitation has also been shown to induce metabolic alterations in heterotrophic bacteria, primarily in respiration and carbon metabolism^26^. In order to acquire this essential trace metal, bacteria can utilize both ferrous (Fe(II)) and ferric (Fe(III)) iron uptake mechanisms. Ferrous iron uptake systems (i.e. Feo) are limited in marine systems, as this metal species is prevalent only in highly acidic, reducing, or anaerobic conditions^27, 28^. Conversely, ferric iron uptake systems are common in marine organisms and comprise three different mechanisms: siderophore-mediated uptake, direct iron reduction, and uptake via transport proteins. Siderophores are small molecules with a high iron (III) binding affinity, and their biosynthetic genes are among the most prevalent in ocean microbial metagenomes, exhibiting high phylogenomic distribution^29^. The first marine siderophores, bisucaberin and anguibactin, were characterized in the late 1980s^30, 31^. Since then, multiple siderophores have been identified in seawater from metagenomic, metatranscriptomic and metabolomic studies^32–40^. They collectively account for at least 0.2-10% of the oceanic dissolved iron pool, evidencing the role that marine microbes play in iron cycling and speciation^32, 36, 37^. Amphibactins and petrobactin are of particular interest, as they are some of the most abundant and widely distributed siderophores in the ocean^36, 38, 41^. Structurally, siderophores are classified by the specific substructures they use to chelate iron. The most common substructures are catecholate, hydroxamate and α-hydroxycarboxylate (**Figure S1**). Additional substructures include arylthiazoli(di)ne, aryloxazoli(di)ne, and diazeniumdiolate^42, 43^. However, siderophores often contain multiples of these, thereby being classified as mixed siderophores.^44^.

The work presented in this manuscript is centered around discovery of siderophores and other mechanisms of iron acquisition in *Microbulbifer* sp. CNSA002 bacteria^45^. With this work, we expand upon the known *Microbulbifer* natural product repertoire by characterizing the first siderophore systems in this genus and describe unprecedented longer chain length acylation on the central spermidine moiety of petrobactin, which is one of the most abundant and widely distributed siderophore characterized from marine environment. These siderophores were only produced in monoculture and not in mixed culture with the marine pathogen, *V. coralliilyticus*. We posit that deciphering how these siderophores shape *Microbulbifer*’s interspecies interactions and the regulation of their production will contribute to the understanding of their function in microbial chemical ecology.

## RESULTS AND DISCUSSION

### Siderophore production by *Microbulbifer* sp. CNSA002

To investigate the ability of *Microbulbifer* sp. CNSA002 to obtain iron, we sequenced its genome and used the MiGA (Microbial Genomes Atlas)^46^ platform to confirm its quality (95.5%) and completeness (100%). We analyzed the genome for the presence of iron chelating proteins, and biosynthetic gene clusters (BGCs) for siderophore biosynthesis. Genome analysis revealed the presence of multiple open reading frames encoding iron-binding and transport proteins (**Table S1**). This analysis showed that the strain CNSA002 has the genetic potential to import siderophores produced by other microorganisms, also called xenosiderophores, via TonB siderophore receptors. Additionally, two BGCs of interest were identified using antiSMASH platform (**Figure 1A**).^47^ One BGC was predicted to encode a nonribosomal peptide synthase (NRPS) metallophore, presenting low similarity (20%) to enantio-pyochelin. The other BGC was predicted to encode an NRPS-independent siderophore (NIS). Despite showing no similarity to known clusters, it was classified as such due to the presence of two *iucA*/*iucC* (iron uptake chelate) genes. To investigate the production of siderophores experimentally, we employed a modified chrome azurol S (O-CAS) agar assay^48^ (**Figure 1B**). The color change from blue to orange observed in and around the *Microbulbifer* sp. CNSA002 colony demonstrated its ability to chelate iron. To assess the presence of siderophores, we fractionated the culture supernatant into the small molecule and macromolecule fraction using a 3 kDa molecular weight cut off (MWCO) filter. The color change in both the concentrated flow through and retentate indicated the production of iron-chelating compounds in addition to iron-chelating proteins (**Figure 1C**).

**Figure 1.**
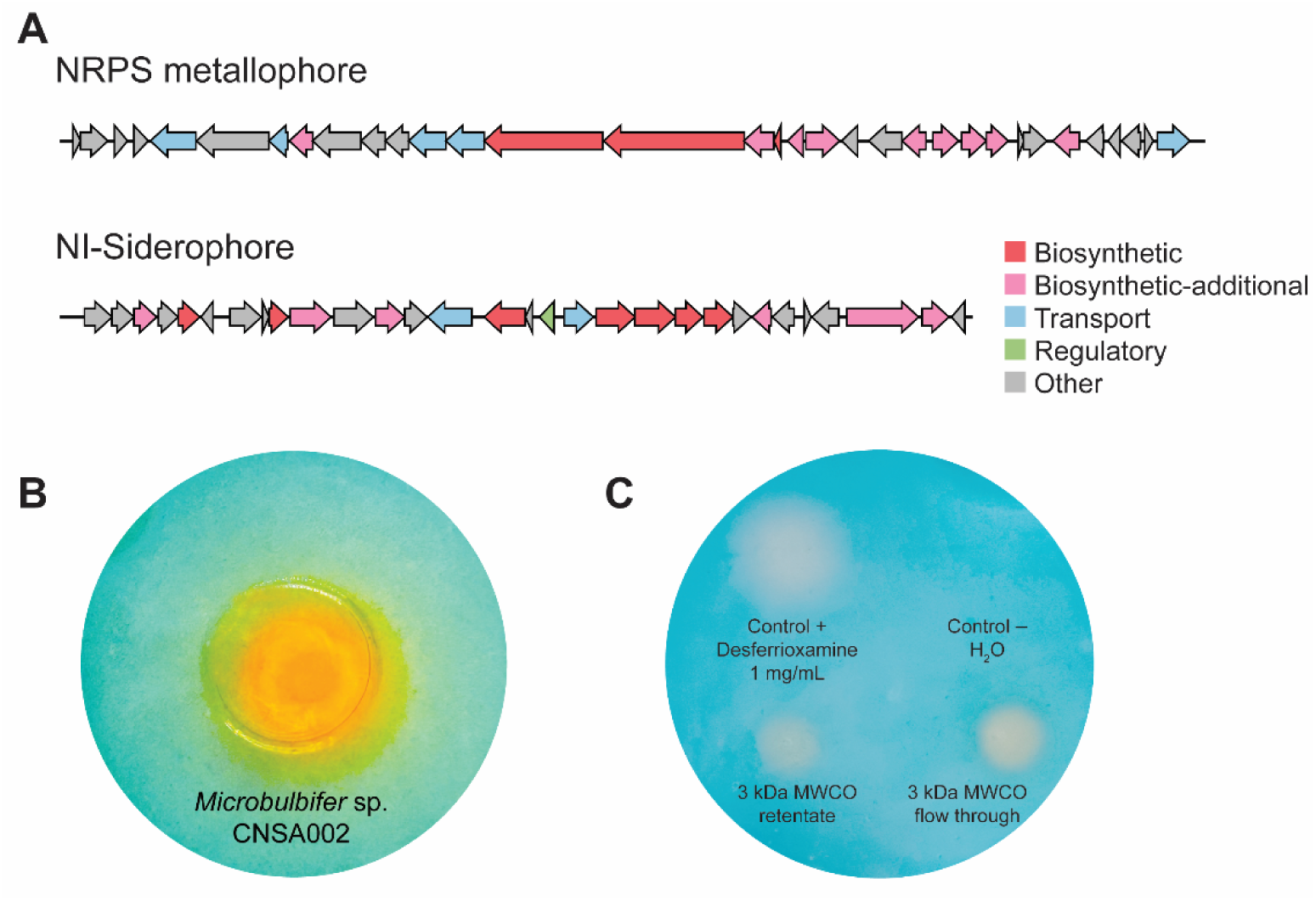
Production of siderophores by *Microbulbifer* sp. CNSA002. **A)** Siderophore BGC candidates identified by antiSMASH on whole genome sequence data for *Microbulbifer* sp. CNSA002. **B)** O-CAS agar assay of *Microbulbifer* sp. CNSA002 colony. **C)** O-CAS agar assay of *Microbulbifer* sp. CNSA002 protein (3 kDa MWCO filter retentate) and small molecule (3 kDa MWCO filter flow through) fractions.

### Mass spectrometry query language-based discovery of bulbichelins

To identify the siderophores produced by *Microbulbifer* sp. CNSA002, we cultured this strain in complex media with and without the supplementation of ethylenediamine-*N*,*N’*-diacetic acid (EDDA). The metabolites were extracted using three extraction methods to capture a wide array of compounds: liquid-liquid extraction (LLE), solid phase extraction (SPE) and solid-liquid extraction (SLE) followed by tandem mass spectrometry (MS) data acquisition. The tandem MS spectra were used to generate a feature-based molecular network (FBMN) in the global natural products social molecular networking (GNPS)^49^ platform as described in the methods. In this network, the structurally similar features are grouped into clusters of connected nodes by quantifying MS^2^ spectral similarity. This workflow additionally aids in the annotation of known natural products, as it computes the spectral similarity between the input data and the GNPS MS^2^ spectral library, which consists of over 500,000 tandem MS spectra. Despite our search against this considerable database and others such as MarinLit and dictionary of natural products, no matches to any known siderophores were identified.

Therefore, we applied MassQL^50^ to identify iron-binding small molecules. This query uses defined patterns of parent *m/z* (MS^1^) and fragmentation data (MS^2^) such as intensity of fragment ions, adducts, isotopic patterns, and neutral losses, to find specific classes of compounds. The isotopic pattern of Fe, in addition to the mass shift between iron-bound and iron-free ligands, makes siderophore amenable to mass spectrometry identification^51–53^. These two characteristics, easily translatable into MassQL queries, underpin the proposed use case for siderophore search using the tool^54^. However, this query did not result in any matches within our dataset, presumably due to lack of detection of iron-bound siderophores. Therefore, we generated a comprehensive library of siderophore substructures (**Figure 2A**, **Table S2**) and wrote queries to mine our dataset for presence of such substructures (**Table S3**). Siderophore discovery through MS^2^ mining has been previously implemented in MATLAB^55^, however, the approach described herein covers a larger and comprehensive library of substructures and employs a web-browser-based tool^50^. The 30 substructures employed for siderophore mining belong to five different families, namely, α-hydroxycarboxylate, catechol, diazeniumdiolate, hydroxamate and aryl thiazoli(di)ne/oxazoli(di)ne (**Figure 2A, Table S2**). The results from each substructure was analyzed using molecular network in GNPS and subsequently visualized in Cytoscape^56^. Spectral matches to non-siderophore compounds were subtracted, yielding a network of 1812 nodes. In this network we identified three clusters of interest (**Figure 2B**). Notably, cluster 1 was uniquely detected under iron-limited conditions (EDDA addition), whereas clusters 2 and 3 were detected in rich media (without EDDA) to a greater relative abundance.

**Figure 2.**
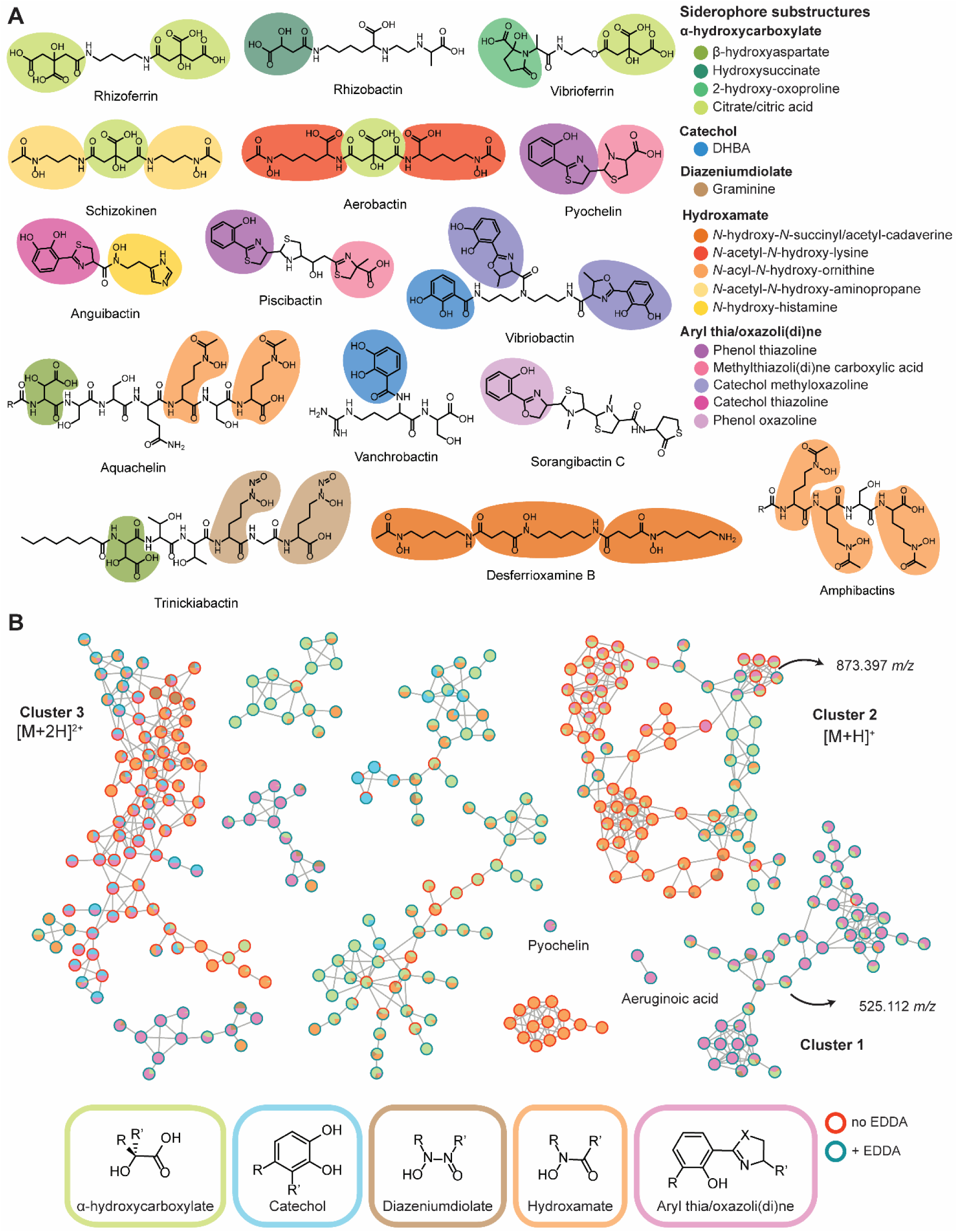
**A)** Representative substructure library for MassQL searches. Siderophore families are grouped by color scheme and specific moieties are represented in different colors. DHBA: dihydroxybenzoic acid. **B)** Representative feature-based molecular network of MassQL results. Clusters of interest and features at *m/z* 525.112 and 873.397 are labeled. Siderophore families are shown, R and R’ represent diverse aryl or alkyl groups, X= S or O. Node pie charts represent the proportion of query hits for each family and border color represent culturing conditions (with or without EDDA).

### Cluster 1: NRPS metallophore

We predicted the molecular formula for the feature at 525.112 *m/z* in this cluster to be C_22_H_29_N_4_O S ^+^.This cluster had no spectral hits to the GNPS library, but matches to the phenol thiazoline, 2-hydroxy-5-oxoproline, *N*-formyl-*N*-hydroxy-ornithine and β-hydroxyasparagine substructures were identified using MassQL (**Figure 3A**). We conducted database searches for the phenol thiazoline substructure and the corresponding neutral mass, however, these searches yielded no results. We then used the MASST^57^ tool to search for the feature of interest against publicly available MS^2^ datasets in the MassIVE repository. MASST is a web-based tool that allows users to search for identical or similar MS^2^ spectra in data repositories, regardless of whether the compound is annotated. This search didn’t result in any analogous spectra either, which prompted us to characterize this compound as a potentially new marine siderophore. The identification of this compound was additionally motivated by a recent phylogenetic inventory, which demonstrated the proclivity of *Microbulbifer* spp. to produce iron chelators, and the lack of siderophores described from the genus^10^.

**Figure 3.**
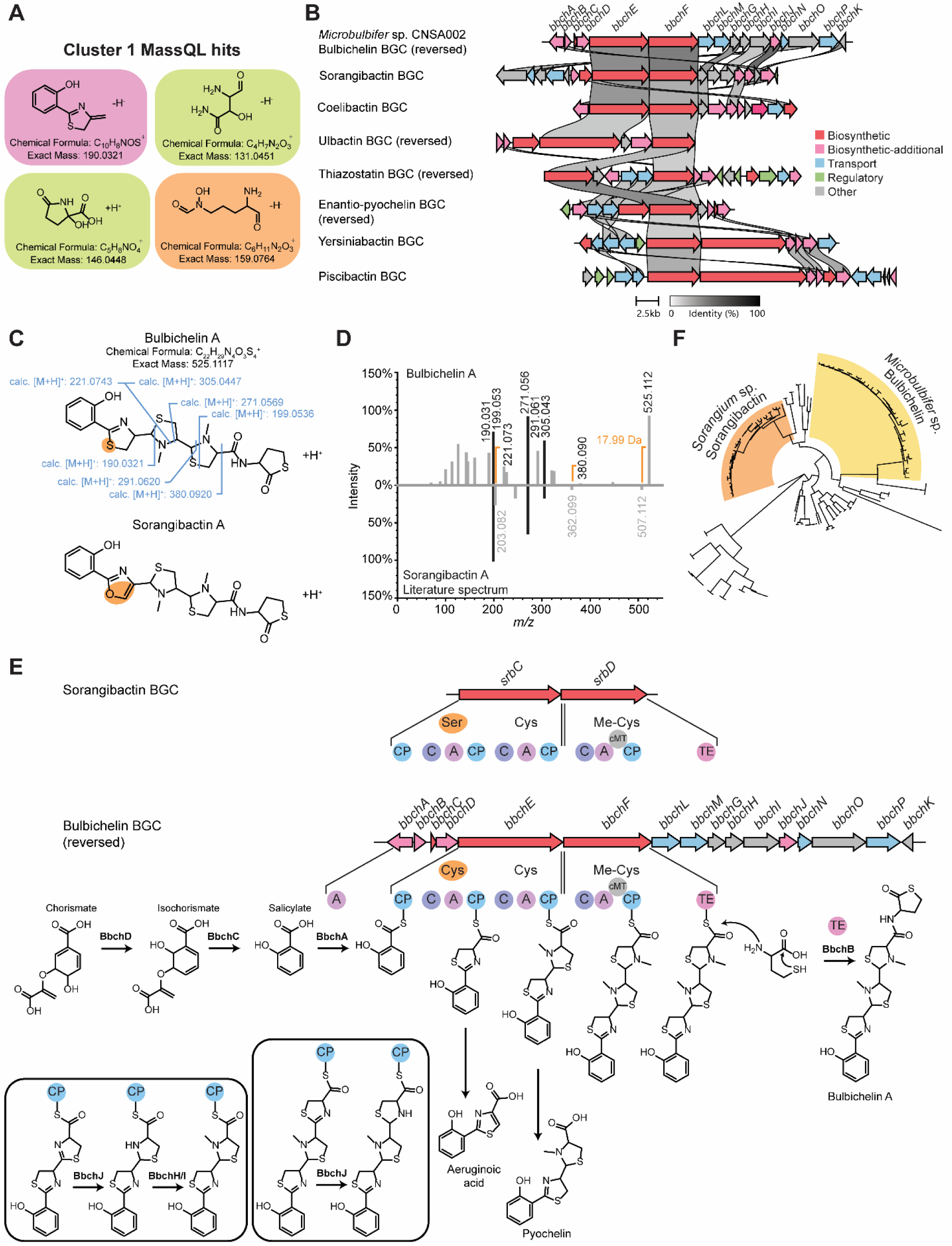
**A)** Most abundant MassQL matches identified in cluster 1: phenol thiazoline, 2-hydroxy-5-oxoproline, *N*-formyl-*N*-hydroxy-ornithine and β-hydroxyasparagine **B)** Synteny analysis of aryl thia/oxazoli(di)ne siderophore BGCs. **C)** Structure of bulbichelin A and sorangibactin A. Fragmentation is shown for bulbichelin A and structural differences are highlighted in orange. **D)** Mirror plot comparing fragmentation spectra of bulbichelin A and sorangibactin A. NMR data is provided as supplemental information. **E)** NRPS module architecture and adenylation domain prediction comparison for bulbichelin (*bbch*) and sorangibactin (*srb*) and proposed bulbichelin biosynthesis. **F)** Phylogenetic tree depicting BbchE homologous proteins. *Microbulbifer* sp. and *Sorangium* sp. clades are pointed out.

To inform the structural characterization of compounds in cluster 1, we performed a synteny analysis of the NRPS metallophore BGC from *Microbulbifer* sp. CNSA002 and aryl thia/oxazoli(di)ne siderophore BGCs (**Figure 3B and S2**) using CAGECAT^58^ clinker tool, with the results showing good alignment of the core biosynthetic genes. AntiSMASH^47^ had predicted the most similar known cluster to be enantio-pyochelin, however, the synteny and phylogenetic analyses revealed a higher similarity to the sorangibactin BGC^59^ (**Figure 3B**). Based on BGC and MS^2^ spectral similarity between sorangibactin and the 525.112 *m/z* produced by our *Microbulbifer* sp. CNSA002, we proposed its structure and named it bulbichelin A (**Figure 3C**). Spectral analysis supports the similarity between bulbichelin A and sorangibactin A, with multiple fragments coinciding for both compounds (**Figure 3D**). Notably, two fragment peaks (221.073 and 380.093 *m/z*) exhibit a mass shift of 17.99 Da when compared to the sorangibactin spectrum. Additionally, substrate prediction in antiSMASH for the adenylation domains in these two BGCs (sorangibactin BGC, *srb*, and bulbichelin BGC, *bbch*) differ in the incorporation of a cysteine in bulbichelin in place of serine in sorangibactin (incorporated by BbchE and SrbC, respectively) (**Figure 3E**). This biosynthetic discrepancy supports the structural and spectral differences observed. Indeed, this Δ*m/z* corresponds to the incorporation of O in place of S (-15.9771 Da) and loss of one degree of unsaturation (-2.014 Da). The discovery of sorangibactin brought forward the unprecedented incorporation of a thiolactone at the C-terminus, which is also present in our bulbichelin with an additional substitution of oxazole in sorangibactin to thiazoline in bulbichelin.

A phylogenetic tree was constructed incorporating the 100 most similar sequences to BbchE (**Figure 3F**). The tree suggests a close relationship between the *Sorangium* sp. and *Microbulbifer* sp. clades, underscoring the similarity of their biosynthetic products. The phylogenetic tree also revealed anthrochelin (*ant*) as an additional BGC of interest, yielding biosynthetic products that are structurally related to sorangibactins^60^. Furthermore, the tree highlights the prevalence of BbchE in the *Microbulbifer* genus, being identified in 34 publicly available genomes, in addition to *Microbulbifer* sp. CNSA002. This is consistent with a recent phylogenetic inventory, which identified this gene cluster to be present in 38 *Microbulbifer* sp. genomes (genomic species GS-I, GS-II, GS-IV (associated with *M. variabilis* ATCC 700307) and some GS-III)^10^.

Sequence homology and genomic analysis have aided in constructing a proposed biosynthesis for bulbichelin, containing a N-terminal phenol, a thiazoline, two *N*-methylthiazolidines, and the C-terminal γ-thiolactone moiety (**Figure 3E**). This BGC is composed of 16 annotated proteins involved in biosynthesis and transport (**Table S4**). Biosynthesis is initiated by salicylate formation from chorismate, proposed to be catalyzed by BbchD and BbchC. These enzymes are predicted to be an isochorismate synthase and lyase, respectively. In anthrochelin, these reactions are carried out by AntN and AntM, respectively. This process differs from the proposed biosynthesis for sorangibactin, where salicylate is synthesized from chorismate via a one-step reaction, catalyzed by the salicylate synthase SrbB^59^. The salicylate moiety is then adenylated by BbchA and loaded onto the first BbchE carrier protein. BbchE and BbchF comprise three adenylation and heterocyclization domains, predicted to use cysteine as substrates for chain elongation. Chain release is then carried out by a thioesterase (TE) domain. Multiple sequence alignment, comparing the TE domains in SrbD, AntB and BbchF revealed a conserved use of the rare catalytic cysteine, instead of serine, as the nucleophile in the enzyme active site (**Figure S3**). This TE likely uses homocysteine as an intermolecular nucleophile, as has been observed for select TEs^59–61^. The intermolecular offloading reaction yields a linear product, thus, thiolactone formation from homocysteine remained unclear in sorangibactin biosynthesis. We hypothesize that a stand-alone TE (BbchB) carries out the thiolactonization of the homocysteine (**Figure 3E**). A homologue of this enzyme is present in the *ant* BGC (AntL), however, no homologues were identified in the *srb* BGC. Stand-alone thioesterases (type II TEs) are usually not involved in biosynthesis, performing an editing role instead by removing unsuitable substrates that block the assembly line^62^. Still, some type II TEs perform non-canonical functions^63^. For instance, SncF catalyzes a lactam-forming cyclization to yield the product spinactin A^64^. A phylogenetic analysis was carried out, including TEs with known crystal structure in the ThYme database^65^, in addition to SncF, BbchB and AntL. The phylogenetic tree shows clustering of these three TEs within the TE 18 family (**Figure S4**), which supports our proposal of thiolactonization being carried out by BbchB; however, *in vitro* experiments are required to validate this hypothesis.

In addition to these core enzymes, BbchH-BbchJ are tailoring enzymes, working *in trans* as oxidoreductases and methyltransferases to yield bulbichelin A (**Table S4**). Notably, the known siderophores pyochelin and aeruginoic acid were also detected upon iron limitation (**Figure 2B**). Given the similarity of these compounds to bulbichelin, and the lower intensity of these ions, we hypothesize that these are shunt products of the bulbichelin BGC, which is consistent with the proposed biosynthesis (**Figure 3E**).

The similarity between the bulbichelin, anthrochelin and sorangibactin BGCs, revealed through synteny and phylogenetic analyses, informed the dereplication of bulbichelin analogs (**Figure 4A and S5, Table 1**). Bulbichelin B bears resemblance to sorangibactin D and anthrochelin B, harboring a carboxylic acid on the C-terminal end and suggesting a hydrolytic release from BbchF TE domain. Bulbichelin C, on the other hand, bears similarity to sorangibactin B and anthrochelin C, where the linear homocysteine suggests it is an intermediate product, prior to the formation of thiolactone. This linear homocysteine could also stem from hydrolytic ring opening due to chromatographic conditions. Interestingly, this thiol group can form disulfide bonds through oxidation, as observed in bisanthrochelin. Bulbichelin C is also subject to this reaction resulting in dimeric bisbulbichelin. Given the proclivity to both ring opening and dimerization during the isolation process, bisbulbichelin was the most amenable to structural confirmation via nuclear magnetic resonance (**Figure S6-10, Table S5**).

**Figure 4.**
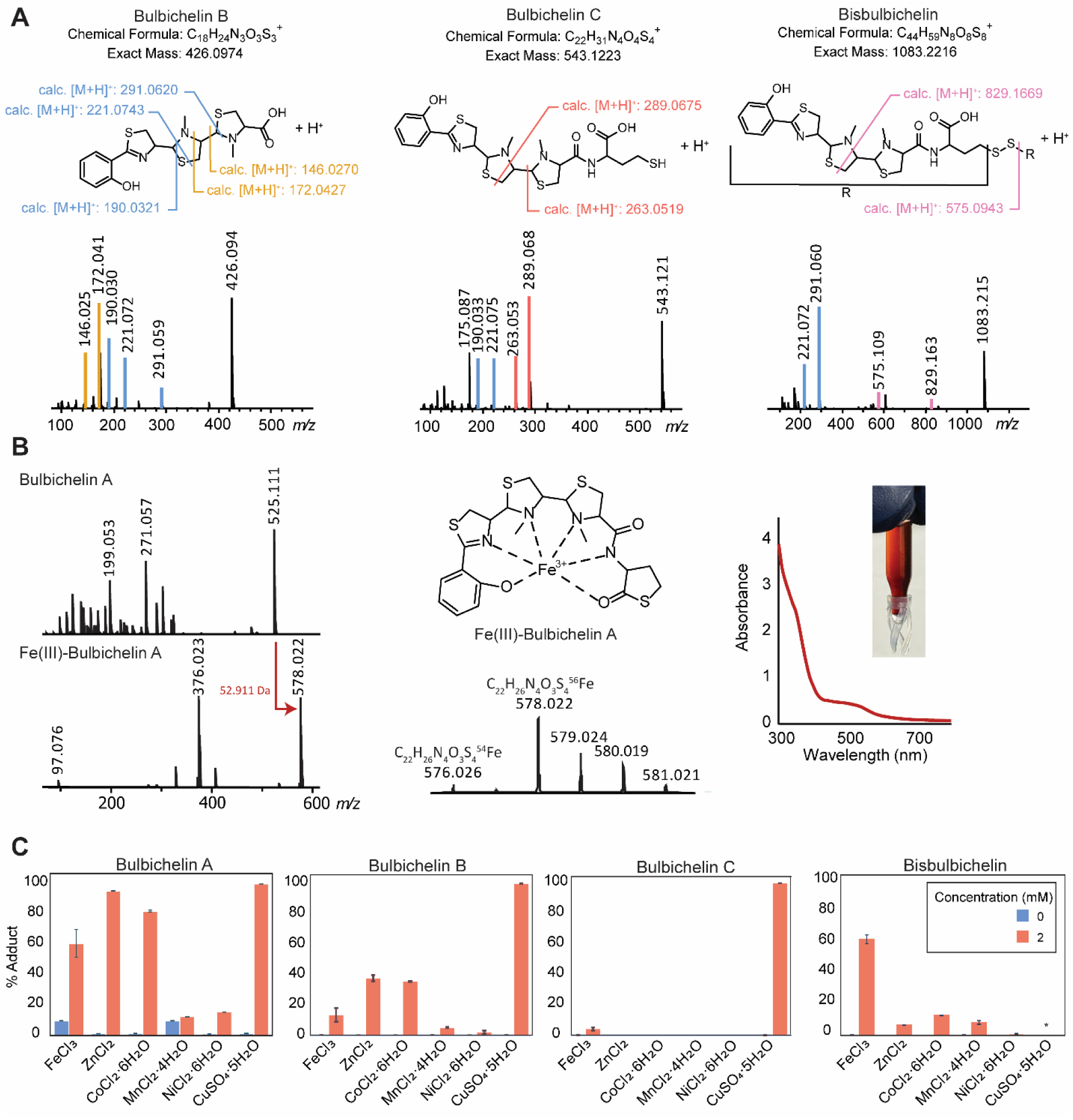
**A)** Bulbichelin analogs MS^2^ fragmentation **B**) Fe(III)-Bulbichelin A chelate spectra, isotopic pattern and UV-Vis spectrum **C)** Bulbichelin analogs metal binding in direct infusion experiments. Binding percentage (% adduct) is defined as the peak area of the metal adduct divided by the total peak area of the metal adduct and the [M+H]^+^ adduct in each infusion run. N=3 for each water infusion (control 0 mM; blue bars) and metal infusion (2 mM; red bars). Several copper adducts are indicated as 100% because proton-bound peaks were either not found or were below the limit of detection, suggesting complete conversion. The asterisk indicates that bisbulbichelin exhibits the adduct bound to two copper ions along with potential reactivity with copper.

**Table 1.**
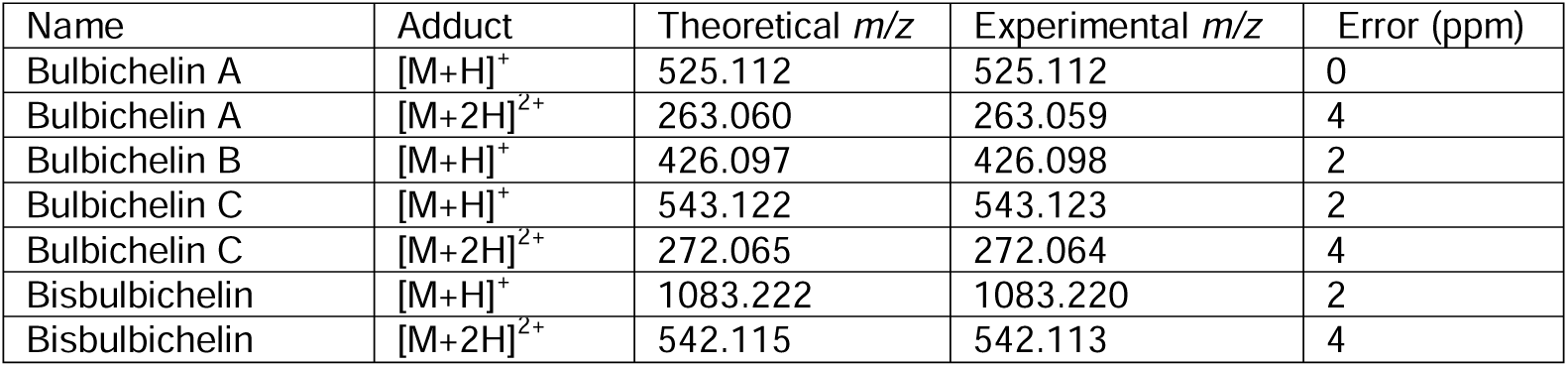
Bulbichelin analogs detected in *Microbulbifer* sp. CNSA002 iron-limited culture.

### Metal binding by bulbichelins

The metal binding properties of bulbichelins were determined using mass spectrometry. Direct metal infusions, allowed for the detection of iron-bound bulbichelins, supporting their potential role as siderophores (**Figure 4B and C**). These iron complexes exhibit the characteristic Δ*m/z* of 52.911 Da, stemming from the [M+Fe-2H]^+^ adduct, and distinct isotopic pattern reflecting the stable isotope ratio between ^54^Fe and ^56^Fe (**Figure 4B and S11**). Bulbichelin A exhibited the highest iron-binding percentage, while analogs B, C, and bisbulbichelin exhibited lower iron-binding percentages (**Figure 4C**). Here, the binding percentage is defined as the metal-adduct peak area divided by the total peak area of the metal adduct plus the [M+H]^+^ adduct in each infusion run. Notably, the chelates display a red-orange color due to strong ligand-to-metal charge transfer bands in the UV-Vis region (HPLC vial insert in **Figure 4B**). Additional direct metal infusion experiments were carried out with a panel of transition metals, given the demonstrated metallophore capabilities of the aryl thiazoli(di)ne/oxazoli(di)ne family. For instance, yersiniabactin is known to chelate zinc, copper, cobalt, nickel, chromium and gallium in addition to iron^66–70^. Similarly, micacocidin can also bind zinc, manganese, copper and gallium^71, 72^; likewise, pyochelin has been shown to chelate a myriad of metals, including the ones previously mentioned^73^. In these native infusion mass spectrometry experiments, bulbichelins also demonstrate metal-binding promiscuity (**Figure 4C**). Bublichelins A, B, and C exhibit appreciable copper binding, and bisbulbichelin shows binding to two copper ions (**Figure S12**). Furthermore, bulbichelin A exhibited high binding percentages for zinc- and cobalt. Bulbichelin B also binds these two metals, though perhaps less readily based on lower binding percentages (**Figure 4C**), while bulbichelin C and bisbulbichelin show low to negligible binding of these two metals. Interestingly, bulbichelins A and B show higher binding ratios of zinc, cobalt, and copper as compared to iron. Finally, while bulbichelin A and bisbulbichelin demonstrate low nickel and manganese binding, any nickel or manganese binding to bulbichelins B and C is below the limit of detection for these experiments, suggesting minimal binding (**Figure 4C**). Thus, bulbichelins may serve as broad spectrum metal binders, a beneficial trait in the dilute marine environment.

### Clusters 2 and 3: NI-siderophore

Clusters 2 and 3 represent [M+H]^+^ and [M+2H]^2+^ adducts of the same compounds, and were therefore analyzed in parallel. These clusters had MassQL hits belonging to dihydroxybenzoic acid, graminine, catechol methyloxazoline, citrate and several hydroxamate queries (**Figure 5A**). The presence of catechol in these features was additionally evidenced by the characteristic 3,4-dihydroxybenzoic acid UV absorbance ratio at wavelengths 294 nm and 255 nm^74–76^ (**Figure 5B**). No nodes in these clusters had matches to the GNPS spectral library and database searches for these substructures and *m/z* did not yield any candidate siderophores.

**Figure 5.**
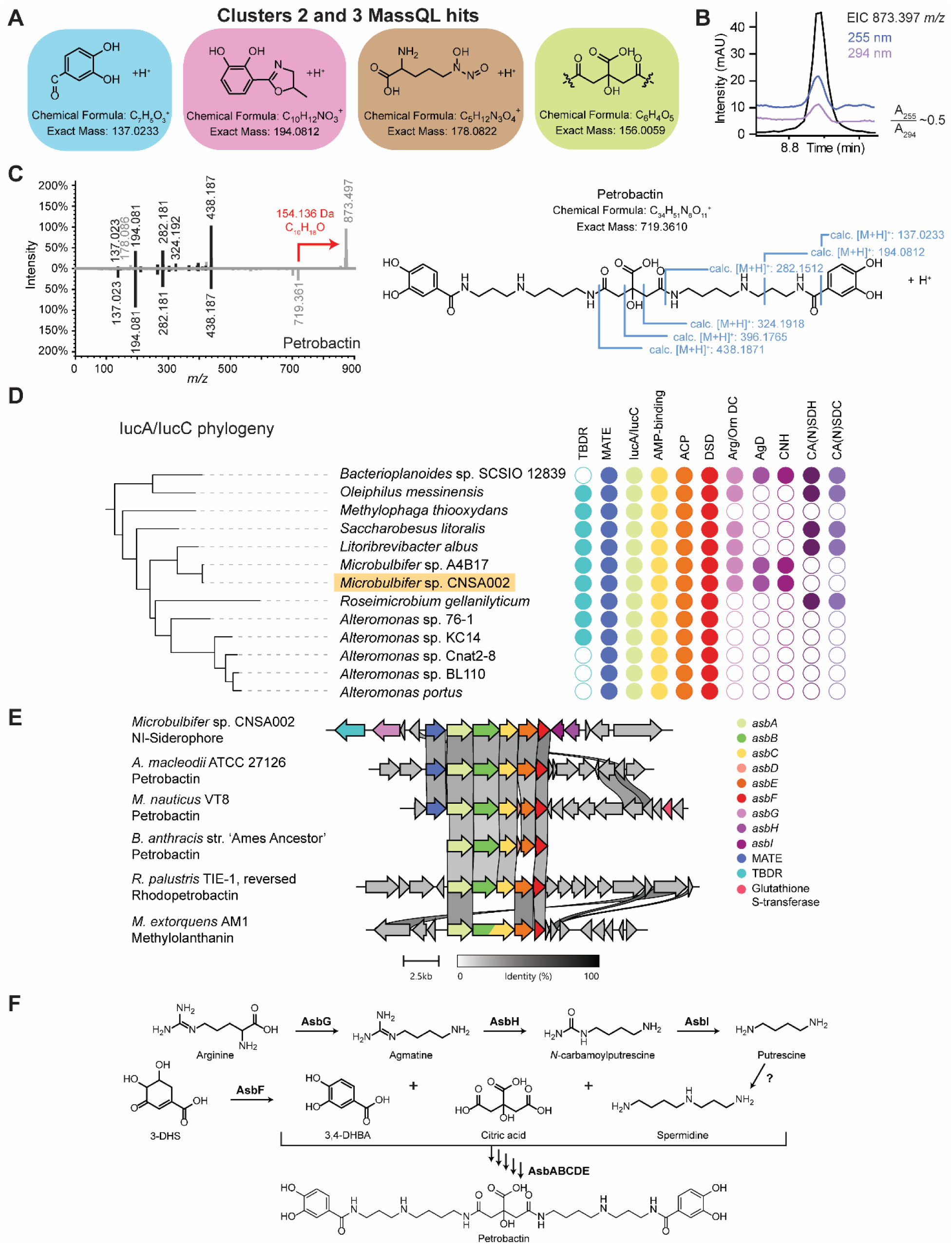
**A)** Most abundant MassQL matches identified in clusters 2 and 3: dihydroxybenzoic acid, catechol methyloxazoline, graminine, and citrate. **B)** Extracted ion chromatogram for feature of interest at 873.397 *m/z* and UV chromatograms for wavelengths 294 nm and 255 nm. **C)** Mirror plot comparison of MS^2^ fragmentation for feature of interest at 873.397 *m/z* and the siderophore petrobactin. The Δ*m/z* between the parent ions is 154.136 Da. Petrobactin MS^2^ fragments are annotated. **D)** IucA/IucC protein phylogeny and co-association analysis. Circles denote presence/absence in gene neighborhood. TBDR: TonB-dependent receptor; MATE: MATE (multidrug and toxic compound extrusion) family efflux transporter; IucA/IucC: IucA/IucC siderophore biosynthesis; ACP: Aryl-carrier protein; DSD: 3-dehydroshikimate dehydratase; Arg/Orn DC: arginine/ornithine decarboxylase; AgD: agmatine deiminase; CNH: carbon-nitrogen hydrolase; CA(N)SDH: carboxy(nor)spermidine dehydrogenase; CA(N)SDC: carboxy(nor)spermidine decarboxylase. **E)** Synteny analysis comparing the NI-siderophore BGC from *Microbulbifer* sp. CNSA002, siderophore clusters from petrobactin-producing bacteria and the published petrobactin BGC from *B. anthracis*. BGCs from the structurally related compounds rhodopetrobactin and methylolanthanin are also shown. **F)** Petrobactin biosynthesis including AsbG: arginine decarboxylase, AsbH: agmatine deiminase, AsbI: *N*-carbamoylputrescine amidase. 3-DHS: 3-dehydroshikimate; 3,4-DHBA: 3,4-dihydroxybenzoic acid

We then performed a MASST search for similar spectra in publicly available MS^2^ datasets. One dataset (MSV000084873, **Figure S13**) contained a MassQL query hit to the known NI-siderophore petrobactin. The fragmentation pattern of this metabolite and our features of interest was remarkably similar (**Figure 5C**). Therefore, we hypothesized that these features are structurally related to petrobactin. We then performed a phylogeny and co-association analysis based on the IucA/IucC present in the genome of *Microbulbifer* sp. CNSA002. The *iucA*/*iucC* genes were first described in the biosynthesis of aerobactin^77, 78^ and are now considered as the model biosynthetic genes in NI-siderophores. Biosynthetic pathways encoding for these siderophores often include one to three genes resembling *iucA* or *iucC*, usually responsible for the condensation of citric acid with an amine or alcohol^79^. IucA and IucC homologs have been identified in over 40 bacterial species^80^, while IucA enzymes are also present in fungi and archaea^79^. In this analysis, we identified the clade in which *Microbulbifer* sp. CNSA002 is nested (**Figure 5D**) and used antiSMASH^47^ to predict the NI-siderophores produced by the other organisms in the clade. Here, two BGCs presented similarity to the siderophore roseobactin, although with low confidence (**Table S6**). Roseobactin is a citrate-catechol mixed siderophore produced by *Phaeobacter inhibens* that differs from petrobactin in the orientation of the spermidine moiety, ultimately exchanging the positions of the putrescine and 1,3-diaminopropane subunits^81^ (**Figure S14**). On the other hand, functional annotation of the *Microbulbifer* sp. CNSA002 genome using the RAST (Rapid Annotation using Subsystem Technology) platform^82^ and SEED viewer^83^ (**Table S1**) predicted the presence of anthrachelin biosynthesis genes (*asb* biosynthetic gene cluster). These genes were identified in *Bacillus anthracis*^84^ but later studies determined the product of this BGC to be petrobactin^76^, characterized from the oil-degrading *Marinobacter nauticus* (previously *M. hydrocarbonoclasticus*)^85^. These two annotations therefore support our hypothesis that the observed feature in our study is structurally related to petrobactin.

The petrobactin BGC (*asb*) is particular in its hybrid NIS-NRPS nature similar to the siderophore nocardichelin^86^. The cluster bears two enzymes from the NIS family (AsbA and AsbB), three belonging to the NRPS family (AsbC, AsbD and AsbE) and a dehydroshikimate dehydratase (DSD, AsbF) (**Table S7**). This is one of the few DSDs to be identified in a biosynthetic pathway^87–91^. Homologs of these enzymes co-occur in all the genomes in the CNSA002 clade (**Figure 5D**). Other co-occurring genes include the transport-related TonB-dependent receptor and multidrug and toxic compound extrusion (MATE) family efflux transporter. We additionally compared the *asb* BGC from *Microbulbifer* sp. CNSA002 and other known petrobactin producers, using the CAGECAT^58^ clinker tool. These results show good alignment of the biosynthetic gene clusters compared (**Figure 5E**). Interestingly, the genomic neighborhoods also included polyamine biosynthetic genes. Previous BGC analyses have described petrobactin biosynthesis with spermidine as a substrate^87–89^. We additionally propose the steps involved in the biosynthesis of spermidine based on presence of biosynthetic enzymes in the *Microbulbifer* sp. CNSA002 BGC (**Figure 5E and F**). The *Microbulbifer* sp. CNSA002 *asb* includes an arginine decarboxylase (AsbG), an agmatine deiminase (AsbH) and an *N*-carbamoylputrescine amidase (AsbI), which sequentially produce putrescine (**Figure 5F**). From the co-association analysis, only one bacterium’s BGC in the clade (*Bacterioplanoides* sp. SCSIO 12839) incorporates all the necessary enzymes for spermidine biosynthesis from arginine. The genome of *Microbulbifer* sp. CNSA002 does encode a putative carboxyspermidine dehydrogenase and a carboxyspermidine decarboxylase elsewhere, responsible for spermidine biosynthesis from putrescine.

To identify these putative petrobactin analogs, we compared the fragmentation patterns of our features of interest with the petrobactin MS^2^. Multiple mass shifts between petrobactin and our features of interest corresponded to C_x_H_y_O, congruent with acylation of the petrobactin structure (**Figure 5C and S15**). However, tandem mass spectrometry data of these features was virtually identical to petrobactin’s, the only evident difference being the parent ion and the low intensity y_1_ ion ([M-438.187+H]^+^, **Figure S16**). The lack of incorporation of the acyl modification in the fragment peaks precluded the identification of the acylation site through MS alone.

### Isolation and nuclear magnetic resonance (NMR) structural elucidation

We acquired 1D and 2D NMR data on the purified compounds for structural elucidation (see methods section, **Table S8 and S9, Figure S17-S22**). We performed comparative analysis of NMR data of the acylated and nonacylated petrobactin, facilitating the determination of the acyl position. Spectral analysis revealed an additional carbonyl carbon, indicating a new amide bond within one of the spermidine moieties. Following this observation, we propagated the annotations of additional analogs in the molecular network (**Figure 6A and B**) and annotated the full suite of detected acylated petrobactins (**Figure 6C**, **Table 2**). In this suite, we also identified hydroxyacyl containing petrobactins, a common functionalization present in acyl moieties. Multiple acylated siderophores have been reported from both marine and terrestrial sources, including marinobactins, amphibactins, acinetoferrin and corrugatin to name a few^92–108^. However, the acylation on the central secondary amine observed herein is rare, as most siderophore acylations are terminal (**Figure S23**). The only exception is observed in the siderophore family rhodopetrobactins^74^ (produced by *Rhodopseudomonas palustris* strains) and the metallophore methylolanthanin^109^ (produced by *Methylorubrum extorquens* AM1) (**Figure S23**). However, these compounds incorporate either one or two acetyl groups. Thus, the longer chain acylations (up to C17) described in this work on the central secondary amines are unprecedented in siderophores. Functionalization of this petrobactin moiety with biotin has been synthetically produced to identify uptake proteins for the ferrisiderophore^110, 111^. Thus, modifications in this region are functionally transported inside the cell and are likely to prevent the theft of siderophore by other organisms lacking the specific transporters. Acylated siderophores have shown membrane affinity, partitioning based on acyl tail length, saturation and functionality^112–114^. The amphiphilicity that an acyl tail confers to an otherwise polar headgroup is crucial in aquatic environments, where diffusion of these secondary metabolites represents a high metabolic cost^115^. Acyl tails can act as a tether to the cell membrane^93^, limiting diffusion and facilitating receptor binding^116^. Additionally, diversity in acylating groups creates variability in siderophore hydrophobicity, allowing for siderophores to be either cell-associated or extracellularly secreted. This diversity can generate a gradient where extracellular siderophores transfer iron to membrane associated ones before being imported into the cell^93, 117, 118^. This “shuttle system” is not exclusive to confined siderophore suites, for instance, *Mycobacerium tuberculosis* utilizes a similar strategy where secreted exochelins transfer iron to cell-bound mycobactins^119^. Acylated petrobactins exhibit this membrane partitioning, as longer acyl tail analogs are exclusively detected in the cell extracts, whereas short-chain analogs are only detected in the supernatant extracts (**Table 2**). Other amphiphilic siderophores do not show membrane association, forming self-assembled structures instead under appropriate conditions^92, 120, 121^. Siderophores such as marinobactins, sodachelins and halochelins undergo micelle-to-vesicle transitions, induced by iron chelation. Rather than preventing diffusion, such physical transformations could prevent photodegradation or proteolysis, as has been demonstrated in synthetic lipopeptides^122^.

**Figure 6.**
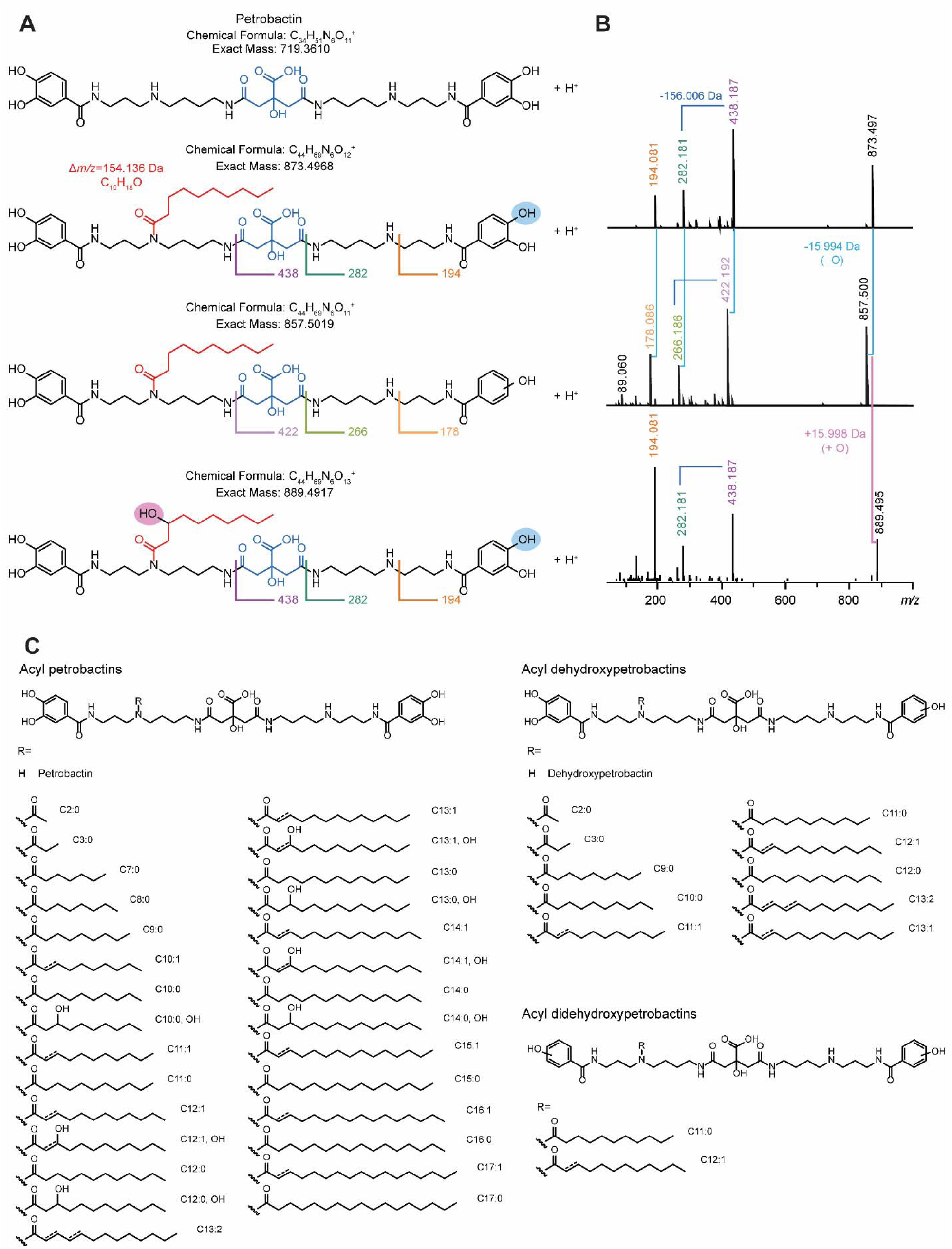
**A)** Representative structure of acyl petrobactins, acyl dehydroxypetrobactins and hydroxyacyl petrobactins, acyl tails are shown in red with the corresponding mass shift. MS^2^ fragments and hydroxyl functional groups are highlighted. **B)** MS^2^ comparison between acyl petrobactins, acyl dehydroxypetrobactins and hydroxyacyl petrobactins, mass shifts corresponding to dehydroxylation are shown in light blue. **C)** Suites of acyl petrobactins, acyl dehydroxypetrobactins and acyl didehydroxypetrobactins.

**Table 2.**
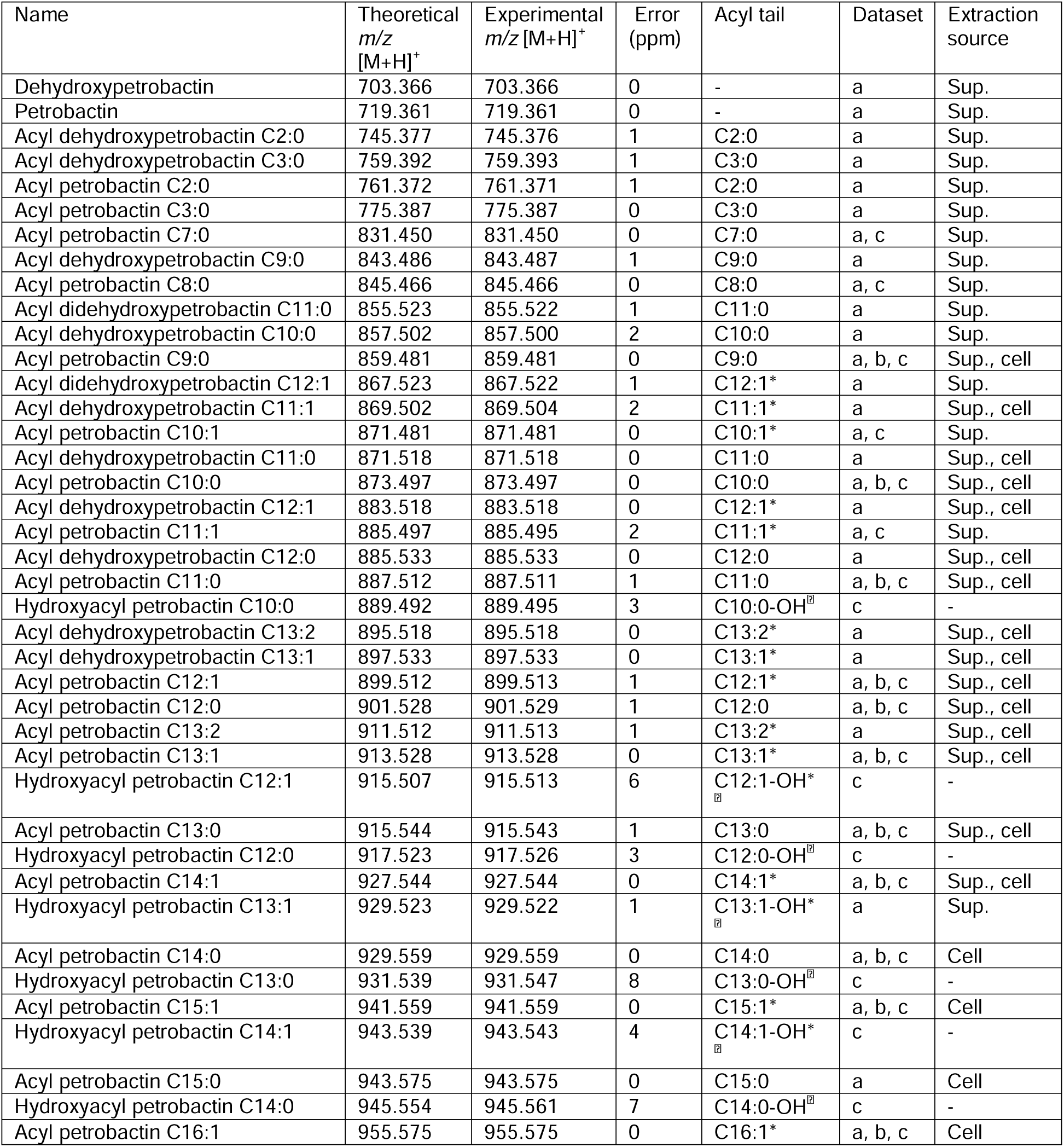
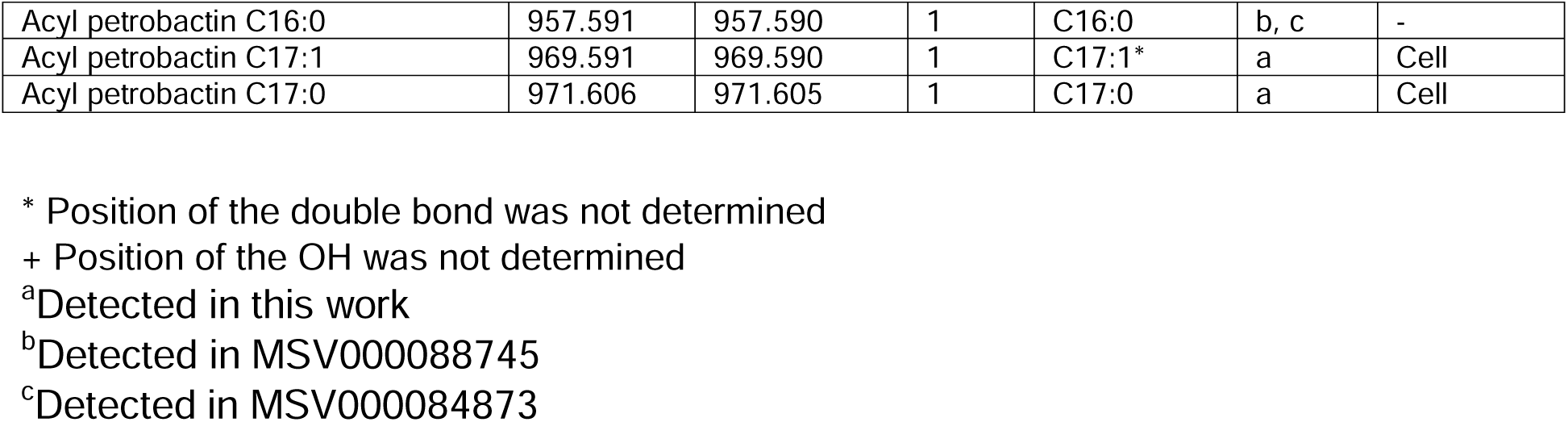
Petrobactin analogs detected in *Microbulbifer* sp. CNSA002 culture and datasets mined with MASST.

Petrobactin is a photoreactive siderophore, where the citrate moiety undergoes an oxidative decarboxylation^85^. This α-hydroxyacid moiety is stable in its metal-free form, but undergoes oxidation after complexation to ferric iron^123^.The petrobactin photoproduct is still able to chelate iron, evidencing the metal-binding ability of the resulting β-ketoenolate^85, 124^. A similar photolability is observed for other siderophores containing central citrates, such as aerobactin and synechobactins^98, 125^. However, we did not detect any acylated petrobactin photoproducts, despite no precautions being taken to prevent photodegradation. The sulfonated petrobactin analogs were also not detected in our work, which is not surprising as our strain does not contain the glutathione S-transferase present in the petrobactin BGC of *M. nauticus* VT8 and ATCC 49840 (**Figure 5E**) ^126, 127^.

### Mass spectrometry-based annotation propagation of petrobactins

Through MS^2^ analysis, we annotated several analogs of acylated petrobactins including the acyl dehydroxypetrobactins and acyl didehydroxypetrobactins, which exchange one or both catechols for a phenol (**Figure 6A and C**). The absence of one hydroxylation renders the dehydroxypetrobactins an asymmetrical molecule, allowing for the detection of both catechol and phenol fragments in MS^2^ (**Figure S15**). On the other hand, didehydroxyopetrobactins are once again symmetrical and only phenol fragments are observed. The incorporation of a phenol instead of a catechol is uncommon for this type of siderophores, as this moiety only contributes one denticity to the overall structure. Siderophores with phenolate moieties usually include them as a part of aryl thia zoli(di)ne /oxazoli(di)ne substructures as discussed above for bulbichelin. Since the *asbF* gene encodes for a 3-dehydroshikimate dehydratase, which converts this substrate into 3,4-dihydroxybenzoic acid^90, 91^, it is unlikely for these analogs to be intermediates in acyl petrobactin synthesis. Instead, these phenol moieties could stem from the ubiquinone pathway, where chorismate is transformed into 4-hydroxybenzoic acid by the chorismate-pyruvate lyase UbiC^128^.

Suites of dehydroxypetrobactins and didehydroxypetrobactins bear one or two fewer iron binding sites compared to acyl petrobactins, losing the preferred hexadenticity for iron chelation. These structural changes have been observed in the aforementioned rhodopetrobactins and the structurally related methylolanthanin. Methylolanthanin incorporates two phenols and is therefore a tetradentate ligand. Despite this denticity, it is still able to chelate iron, but may exhibit a low iron-binding affinity^129^. This compound is primarily described as a lanthanophore, chelating these metals through a complex interplay of iron and lanthanides uptake. This prompts the question of whether petrobactin and its analogs can chelate other metals. Petrobactin is also known to bind boron^130^, a characteristic that is beneficial considering that boron is an essential trace nutrient^131^ and that shows the multiple functionalities exhibited by one compound. This boron-binding ability in siderophores has been observed for vibrioferrin and rhizoferrin as well^130, 132^. Despite being proposed to be antagonistic, as both metals compete for the ligand^131^, the ability to use the same biosynthetic machinery to produce ligands that chelate a wide variety of metals would certainly be advantageous for *Microbulbifer* in the context of nutrient-scarce ocean waters. We confirmed the ability of petrobactin and its acylated analogs to chelate iron through the same O-CAS agar assay (**Figure 7A and 7B**). We tested petrobactin’s affinity for additional transition metals (Zn, Co, Mn, and Ni) through direct metal infusion, however, there was a clear preference for Fe (**Figure 7C**). Acyl petrobactins were also tested, showing the same trend as petrobactin (**Figure S24**).

**Figure 7.**
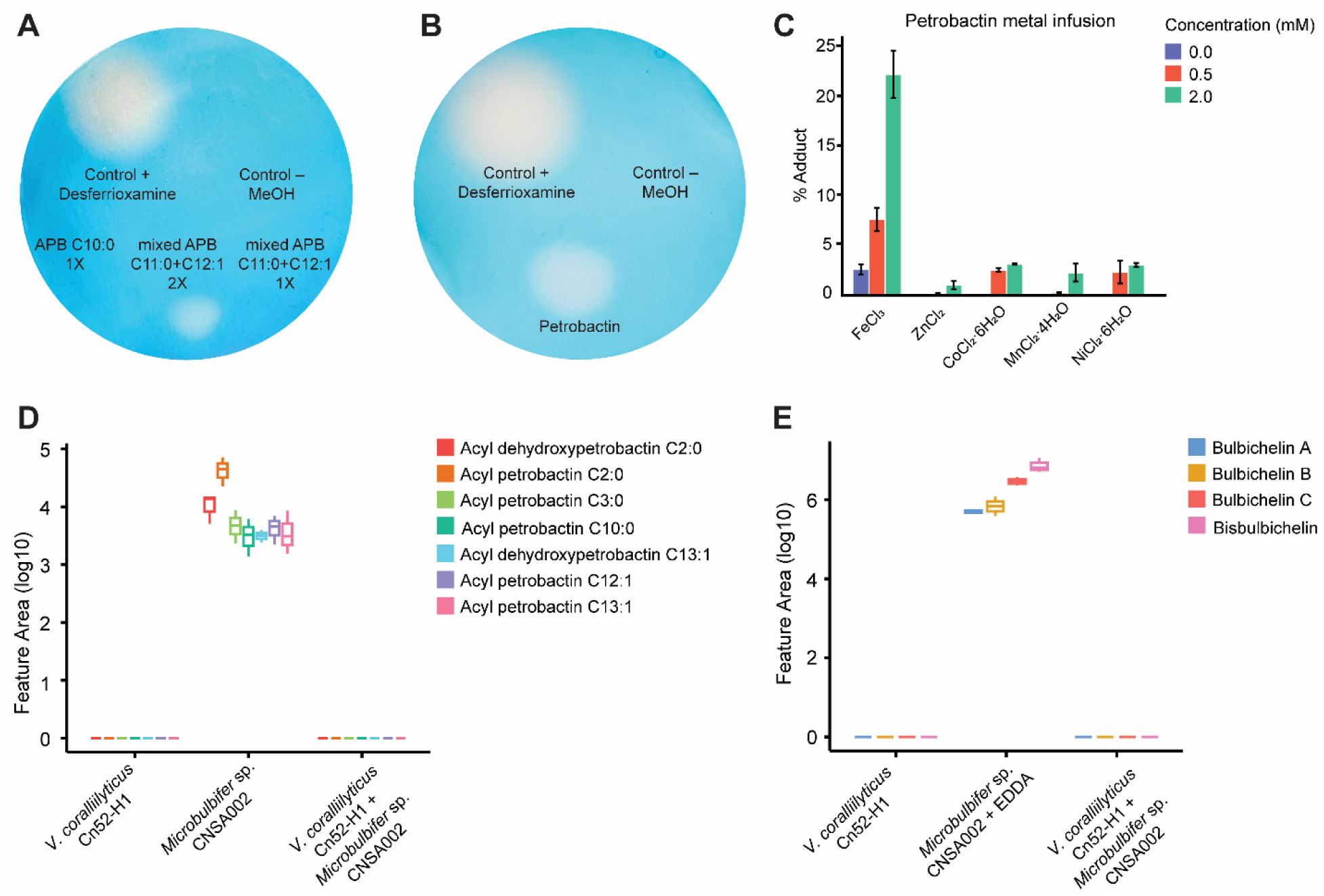
**A)** O-CAS assay for acyl petrobactin (APB) fractions. **B)** O-CAS assay for isolated petrobactin. **C)** Metal binding by petrobactin in direct infusion experiments. Binding percentage (% adduct) is defined as the peak area of the metal adduct divided by the total peak area of the metal adduct and the [M+H]^+^ adduct in each infusion run. N=3 for each metal infusion and metal concentration (0, 2, and 5 mM). **D)** Boxplots of the relative abundances of acylated petrobactins in *Microbulbifer* sp. CNSA002, the pathogen *Vibrio coralliilyticus* Cn52-H1 and their coculture. E) Boxplots of the relative abundances of bulbichelins in *Microbulbifer* sp. CNSA002 grown in iron limited media (EDDA addition), the pathogen *Vibrio coralliilyticus* Cn52-H1 and their coculture.

### Dysregulation of *Microbulbifer* sp. CNSA002 siderophore production upon coculture with the coral pathogen *Vibrio coralliilyticus* Cn52-H1

We had previously reported the ability of *Microbulbifer* sp. CNSA002 to enzymatically degrade the siderophore amphibactin, produced by the marine pathogen *Vibrio coralliilyticus*^15^. Additionally, we demonstrated the constitutive production of this enzyme, underscoring its relevance. This further motivated the identification of *Microbulbifer*’s siderophore(s), in an attempt to unravel the biological implications of this bacterial interaction. We had previously hypothesized that *Microbulbifer* sp. CNSA002 carries out amphibactin degradation to lower the iron binding affinity of exogeneous siderophores. Such a strategy could better position its own iron acquisition mechanisms, however, petrobactin analogs were only detected in the *Microbulbifer* sp. CNSA002 monoculture (**Figure 7D**). Likewise, bulbichelins were not detected in the coculture with *Vibrio coralliilyticus* Cn52-H1 (**Figure 7E**). This observation suggests that *Microbulbifer* sp. CNSA002 does not produce siderophores in coculture with the pathogen and, further, that the presence of *Vibrio*’s amphibactin does not limit iron availability for *Microbulbifer* sp. CNSA002 in the way that EDDA does. This suggests that *Microbulbifer* sp. CNSA002 engages in siderophore piracy, taking advantage of *Vibrio’*s iron acquisition mechanisms. This shift from siderophore production to the utilization of exogenous siderophores has been previously identified in marine bacteria^133^. Amphibactin degradation and the purported decrease in iron affinity of the product could therefore release iron from the chelate. The increase in available iron pool could then downregulate the expression of siderophore BGCs in *Microbulbifer* sp. CNSA002. This strategy would be advantageous for *Microbulbifer*, as the enzyme promiscuity to degrade structurally related NRPS siderophores would grant the theft of iron from siderophores produced by a diversity of organisms. The role of *Microbulbifer* sp. CNSA002 as a siderophore pirate is further supported by the identification of TonB-dependent siderophore receptors and outer membrane receptors for other ferrisiderophores in the genome (coprogen and rhodotorulic acid, **Table S1**). This degradation would also be beneficial given the importance of siderophores in the virulence of pathogenic bacteria. Siderophores can be wielded by pathogens to induce iron starvation in competing microbes, therefore, the degradation of amphibactin could be used as a strategy by *Microbulbifer* to circumvent these efforts by *Vibrio*.

Increased bioavailable iron, often resulting from environmental changes like ocean acidification or anoxic sediment release, significantly impacts microbial ecosystems by shifting metabolic strategies and triggering nutrient competition. Under iron limitation, microorganisms rely on advanced strategies, such as producing siderophores to scavenge iron, directly affecting microbial community composition. The genus *Microbulbifer,* known for degrading complex marine polysaccharides, likely plays a key role in iron cycling, particularly in coastal, iron-limited, or polysaccharide-rich environments. Further research into iron acquisition mechanisms in *Microbulbifer* and their regulation is essential to understand its ecological role in mediating iron availability for other marine organisms or *Microbulbifer*-associated hosts.

### Experimental Section

#### Bacterial culturing

Bacterial culturing, both in mono- and coculture was previously described^15^. Briefly, strains were inoculated in SWB (saltwater broth) with or without the addition of ethylenediamine-*N*,*N’*-diacetic acid (EDDA, Sigma-Aldrich, St. Louis, MO, USA) (300, 500, 750, 1000 µM) to an OD_600_ of 0.05, using a 1:1 ratio for bacterial cocultures. The cultures were incubated at 30 °C, shaking at 200 rpm for 24 h. Harvested cultures were extracted using a variety of techniques: liquid-liquid extraction (LLE) with ethyl acetate (EtOAc), solid phase extraction (SPE) with C18 columns (Thermo Scientific, Waltham, MA) and solid liquid extraction (SLE) with 2:2:1 ethyl acetate:methanol:water (EtOAc:MeOH:H_2_O). Extracts were dried *in vacuo* and stored at -20 °C until data acquisition.

#### Untargeted metabolomics data acquisition and analysis

The dried extracts were resuspended in 300 µL of 100% MeOH containing 1 µM of sulfadimethoxine as an internal standard. Samples were vortexed, sonicated for 10 min and centrifuged at 16,160 x g for 15 min. The resuspended extracts were analyzed using an Agilent 1290 Infinity II Ultra High-Pressure Liquid Chromatography (UHPLC) system (Agilent Technologies; Santa Clara, CA, USA) with a Kinetex 1.7 μm C18 reversed phase UHPLC column (50 × 2.1 mm) (Phenomenex; Torrance, CA, USA) coupled to an ImpactII ultrahigh resolution Qq-ToF mass spectrometer (Bruker Daltonics, GmbH, Bremen, Germany) equipped with an electrospray ionization (ESI) source. Chromatographic separation was performed with the following mobile phase gradient: 5% solvent B (MeCN, 0.1% (v/v) formic acid) and 95% solvent A (H_2_O, 0.1% (v/v) formic acid) for 3 min, a linear gradient of 5% B–95% B over 17 min, held at 95% B for 3 min, 95% B–5% B in 1 min, and held at 5% B for 1 min, 5% B- 95% B in 1 min, held at 95% B for 2 min, 95% B−5% B in 1 min, and held at 5% B for 2.5 min, at a flow rate of 0.5 mL/min throughout. MS spectra were acquired in positive ionization mode from *m/z* 50 to 2000 Da. An external calibration with ESI-L Low Concentration Tuning Mix (Agilent Technologies) was performed prior to data collection, and hexakis(1H,1H,2H-perfluoroethoxy)phosphazene was used as an internal lock-mass calibrant throughout the run. The eight most intense ions per MS^1^ were selected for fragmentation and MS^2^ data acquisition. A basic stepping function was used to fragment ions at 50% and 125% of the CID calculated for each *m/z* with timing of 50% for each step. The MS/MS active exclusion parameter was set to two, and the active exclusion was released after 30s. The mass of the internal lock-mass calibrant was excluded from the MS^2^ list. UV data was acquired with a UV DAD detector (Agilent Technologies) from 190 nm to 400 nm, with a 2 nm step. Zero offset was set at 5% along a 1000 mAU attenuation. Data was acquired throughout the LC run with a >0.1 min peak width.

Raw data were converted to mzXML format, using vendor proprietary software. Metabolite features were extracted using MZmine 2.53^134^, performing mass detection, chromatogram building, chromatogram deconvolution, isotopic peak grouping, retention time alignment, replicate filtering, duplicate peak removal, and gap filling. The resulting processed data was submitted to the Global Natural Product Social Molecular Network platform (GNPS) to generate a feature-based molecular network (FBMN). The molecular network was generated using the following parameters: fragment ions were removed within a ±17 Da window of the precursor *m/z*, precursor ion and fragment ion mass tolerance were set to 0.02 Da and edges were filtered to have a score above 0.7 and at least 4 matched peaks. Edges were kept if both nodes were present in each other’s top 10 most similar nodes, and molecular families’ maximum size was set to 100. Experimental fragmentation spectra were searched against GNPS’s spectral libraries and filtered in the same way (cosine score above 0.7 and a minimum of 4 matched peaks). The workflows for DEREPLICATOR^135^, DEREPLICATOR+^136^, MolDiscovery^137^ and MS2LDA^138^ were also run on GNPS. The output from MZmine was additionally exported for analysis with SIRIUS^139^ 5.3.6. with CSI:FingerID^140^ and CANOPUS^141^. These tools provide putative annotations, which are confirmed using an MS^2^ spectral comparison with literature reported spectra, in-house spectra, with data acquired on analytical standards, and manual annotation of MS^2^ fragments resulting in level 2 compound annotations. SIRIUS computes putative chemical formulas using MS^1^ and fragmentation trees (based on user uploaded MS^1^ isotopic peaks and MS^2^ fragmentation patterns). CSI: FingerID transforms MS^2^ spectra into predicted structural fingerprints that enable matching to chemical databases. CANOPUS predicts the chemical class of metabolites by utilizing CSI:FingerID’s predicted structural fingerprints. Library and database searches were performed for The Natural Product Atlas^142^, LOTUS^143^, MarinLit^144^, Pubchem and Scifinder. The molecular network was visualized using Cytoscape^56^ v3.9.0 and features present in the blanks were subtracted.

A MassQL search was conducted in GNPS for each of the 30 siderophore substructures (**Table S2**). These queries search for common MS^2^ peaks and neutral losses detected for siderophores including each substructure and delimits the *m/z* search to a 10 ppm window and 20% intensity threshold (**Table S3**). The results from each query were extracted and visualized using the classical networking workflow in GNPS.

Regarding cluster 1, a MASST search was conducted in GNPS for the 525.112 *m/z* ion, however, no hits were retrieved. On the other hand, for clusters 2 and 3, a search was conducted for five parent ions (761.371, 873.497, 887.511, 899.513, and 913.528 *m/z*) and five fragment peaks (194.081, 282.181, 324.192, 396.176, and 438.186 *m/z*) shared by the features of interest. Two datasets (MSV000088745 and MSV000084873) contained features matching our search and they were reanalyzed, generating classical networks in order to identify related features.

#### Fermentation, extraction, and compound purification of bisbulbichelin

*Microbulbifer* sp. CNSA002 was inoculated at an OD_600_ of 0.05 in 2.8 L baffled flasks containing 1 L of marine broth at half concentration of nutrients (MB/2) supplemented with 750 µM EDDA. Cultures were incubated at 30 °C, shaking at 180 rpm for 24 h. Harvested cultures (8 L) were extracted two times with ethyl acetate. Extracts were combined and the solvent was dried *in vacuo* to yield a 0.5 g crude extract.

The crude extract was resuspended in MeOH and fractionated using an Agilent 1260 Infinity III Liquid Chromatography (semipreparative HPLC) system (Agilent Technologies) equipped with an Eclipse XBD-C18 5 μm reversed phase HPLC column (250 × 9.4 mm) (Agilent Technologies). The mobile phases used for chromatographic separation were solvent A: H_2_O, 0.1% (*v/v*) formic acid (FA) and solvent B: MeCN, 0.1% (*v/v*) FA. The gradient was performed with the following mobile phase compositions: a linear gradient from 5% solvent B to 100% solvent B in 25 min, held at 100% B for 10 min, at a flow rate of 2 mL/min throughout. Elution was performed with a UV detector monitoring the run at 198 nm, 250 nm, and 352 nm. Fraction 1 contained a mixture of Fe-bound and apo bulbichelin A, fraction 3 contained a mixture of Fe-bound and apo bulbichelin A, as well as bulbichelin C. Fraction 4 contained bisbulbichelin (compound **1**) (1.0 mg).

Compound **1**: white powder. ^1^H and ^13^C NMR data, see **Table S5**. Positive HRESIMS *m/z* 1083.220 [M+H]^+^ (calculated for C_44_H_59_N_8_O_8_S_8_^+^, 1083.2217)

One-dimensional (1D) and two-dimensional (2D) NMR spectra were recorded on a Bruker AV3 HD 700 MHz NMR, in DMSO-*d*_6_ and calibrated using residual undeuterated solvent as internal reference. The chemical shift (δ) is reported in parts per million (ppm) and the coupling constants (*J* values) are in Hz. NMR data for compound **1** has been deposited to NP-MRD^145^ (ID: NP0352519).

#### Fermentation, extraction, and compound purification of acyl petrobactins

*Microbulbifer* sp. CNSA002 was inoculated at an OD_600_ of 0.05 in 1 L flasks containing 200 mL of SWB. Cultures were incubated at 30 °C, shaking at 200 rpm for 24 h. Harvested cultures (7 L) were centrifuged at 4920 × g for 45 min at 4 °C to separate the cell pellet from supernatant. The cell-free supernatant was mixed with 4% (*w/v*) XAD-16N resin (Sigma-Aldrich, St. Louis, MO, USA) and stirred for 16 h. Afterwards, the supernatant was removed, and the resin was washed with 500 mL of H_2_O and eluted with 500 mL of ethanol (EtOH). The cell pellet was disrupted using a tissue homogenizer (Tissue Master 125, Omni International, Kennesaw, GA, USA) and extracted with 50 mL of 2:2:1 EtOAc:MeOH:H_2_O. The extraction was vortexed every 30 min for 8 h. Both extracts were combined and the solvent was dried *in vacuo* to yield a 5 g crude extract.

The crude extract was resuspended in H_2_O and fractionated *via* reverse phase liquid chromatography (RPLC) using a 10 g C18 column (Thermo Scientific, Waltham, MA). The column was washed with 150 mL of MeCN and equilibrated with 150 mL of H_2_O. The crude was loaded, and the analytes were eluted with 20%, 40% and 100% MeCN to obtain three fractions (1-3) which were dried *in vacuo*. Fraction 1 (20% MeCN) was further fractionated using an Agilent 1260 Infinity II Liquid Chromatography (semipreparative HPLC) system (Agilent Technologies) equipped with a Luna 5 μm C18 reversed phase HPLC column (250 × 10 mm) (Phenomenex). The mobile phases used for chromatographic separation were solvent A: H_2_O, 0.1% (*v/v*) TFA and solvent B: MeCN, 0.1% (*v/v*) trifluoroacetic acid (TFA). The gradient was performed with the following mobile phase compositions: a linear gradient from 5% solvent B to 100% solvent B in 15 min, held at 100% B for 10 min, at a flow rate of 2 mL/min throughout. Elution was performed with a UV detector monitoring the run at 190 nm and 254 nm. Fraction 7 (10.2 mg) contained petrobactin and fraction 14 (17.2 mg) contained a mixture of acylated petrobactins. Fraction 7 was further fractionated using the following mobile phase gradient: a linear gradient from 5% solvent B to 100% solvent B in 25 min, held at 100% B for 10 min, at a flow rate of 2 mL/min throughout. This fractionation yielded 5.6 mg of petrobactin. Fraction 14 was further fractionated using the following mobile phase gradient: a linear gradient from 5% solvent B to 60% solvent B in 80 min, 60% solvent B to 100% solvent B in 5 min, held at 100% B for 15 min, at a flow rate of 2 mL/min throughout. This fractionation yielded compound **2** (F14.14) (1.0 mg) and two mixed fractions: F14.15 and F14.16 (2.6 mg and 1.0 mg).

Compound **2**: yellow, oily. ^1^H and ^13^C NMR data, see **Table S8**. Positive HRESIMS *m/z* 873.499 [M+H]^+^ (calculated for C_44_H_69_N_6_O_12_^+^, 873.4968)

Petrobactin: yellow, oily. ^1^H and ^13^C NMR data, see **Table S9**. Positive HRESIMS *m/z* 719.361 [M+H]^+^ (calculated for C_34_H_51_N_6_O_11_^+^, 719.3610)

1D and 2D NMR spectra were recorded on a Bruker Avance III HD 800 MHz NMR, in DMSO-*d*_6_ and calibrated using residual undeuterated solvent as internal reference. The chemical shift (δ) is reported in parts per million (ppm) and the coupling constants (*J* values) are in Hz. NMR data for compound **2** has been deposited to NP-MRD^145^ (ID: NP0351565).

#### O-CAS agar assay

A modified CAS assay (O-CAS^48^) was overlaid on a colony of *Microbulbifer* sp. CNSA002 grown on SWA. The spent SWB media of *Microbulbifer* sp. CNSA002 was fractionated using a molecular weight cutoff (MWCO) Amicon ultracentrifugation filter (3 kDa, Sigma-Aldrich) and each fraction was tested on an O-CAS agar plate. The iron chelating activity of compound **2** and fractions F14.15 and F14.16 were also tested on an O-CAS plate. A 2.5 µL aliquot of each a 200 mg/mL methanolic solution of compound **2**, 400 mg/mL F14.15 and 200 mg/mL F14.16 were spotted on an O-CAS agar plate^48^. Desferrioxamine mesylate (Sigma-Aldrich) 1 mg/mL was used as a positive control, whereas methanol was spotted as a negative control. The plates were incubated at room temperature overnight before registering the results.

#### CNSA002 whole genome sequencing and analysis

We had previously sequenced the *Microbulbifer* sp. CNSA002 genome using Illumina and Nanopore platforms, and deposited to NCBI under the bioproject PRJNA1173768 (NCBI:txid3373604). The genome was analyzed using the PhylaLite and TypeMat workflows under the MiGA platform^46^. Genome annotation was performed in the RAST platform^82^ using the RASTtk^146^ scheme and browsed with SEED viewer^83^. Additional annotation of biosynthetic gene clusters was performed in the bacterial version of antiSMASH^47^ with a relaxed strictness, and NCBI PGAP^147^. AntiSMASH predicted an NRPS metallophore and an NI-siderophore BGC, whereas RAST-SEED predicted the later to be for anthrachelin biosynthesis, a purported NI-siderophore.

We used the CAGECAT^58^ clinker tool to compare the NRPS siderophore BGC from *Microbulbifer* sp. CNSA002 and known aryl thiazoli(di)ne/oxazoli(di)ne siderophore BGCs from MIBiG^148^: sorangibactin (BGC0002912), coelibactin (BGC0000324), ulbactin (BGC0002472), thiazostatin (BGC0001801), enantio-pyochelin (BGC0002475), yersiniabactin (BGC0001055), piscibactin (BGC0002533). *Microbulbifer* sp. CNSA002 NRPS BGC contained two core biosynthetic genes, *bbchE* and *bbchF.* Homologous proteins to BbchE were searched for using the blastp function in NCBI’s BLAST^149^. The 100 most similar sequences were downloaded and aligned using ClustalW^150^ in MEGA11^151^. The aligned sequences were used to generate a phylogenetic tree using the maximum likelihood method and JTT matrix-based model^152^. The tree was visualized in iTOL^153^. BbchF TE domain was compared through multiple sequence alignment with other TE domains, including the sorangibactin and coelibactin thioesterases using MUSCLE^154^ in MEGA11. Thioesterase phylogenetic analysis was performed by downloading sequences of TEs with known protein crystal structure in the ThYme database^65^ (as of February 2026). Sequences for BbchB, AntL and SncF were also added to the analysis. All sequences were aligned using MUSCLE^154^, and a phylogenetic tree was generated in MEGA11 using a JTT matrix-based model^152^. Family nomenclature was assigned based on the ThYme database.

The NIS BGC contained two *iucA/iucC* genes which were input into EFI-EST^155, 156^ (enzyme function initiative-enzyme similarity tool) for analysis using the default parameters. The SSN alignment score threshold was set to 40%. The representative SSN (40% identity) containing 216 nodes was exported and visualized in Cytoscape. The corresponding table was exported as a csv and the accession numbers were used to obtain the RefSeq sequences using the UniProt^157^ ID Map. Protein sequences were retrieved using the NCBI Batch Entrez, non-redundant sequences (139) were aligned using ClustalW^150^ in the CIPRES science gateway^158^. A tree was constructed using the RaxML-HPC Blackbox^159^ workflow using default parameters. The program determined 350 bootstraps to be sufficient, and the output tree was visualized in iTOL^153^. The clade including *Microbulbifer* sp. CNSA002 was identified and selected for further analysis. In order to identify co-occurring genes, the representative SSN was submitted to the enzyme function initiative genome neighborhood tool (EFI-GNT)^156^. The neighborhood size was set to 15 and the minimal co-occurrence limit was set to 20%. The results were visualized in Cytoscape and the respective table was exported. The IucA/IucC entries belonging to the clade of interest were identified and visualized on the genome neighborhood diagram, where co-occurring Pfam families were identified. Protein co-occurrence data was converted into binary annotation and mapped onto the tree using iTOL.

Given the similarity to the petrobactin BGC and the presence of a petrobactin feature in the MSV000084873 dataset with a very similar fragmentation pattern, we used the CAGECAT^58^ clinker tool to compare the NI-siderophore BGC from *Microbulbifer* sp. CNSA002 and published petrobactin BGCs from different bacteria. *B. anthracis* Ames strain petrobactin cluster was obtained from MIBiG^148^ (BGC0000942). The biosynthetic gene clusters for *A. macleodii* ATCC 27126 (NCBI:txid529120) and *M. nauticus* VT8 (NCBI:txid351348, heterotypic synonym *M. hydrocarbonoclasticus*) were obtained by analyzing the whole genome sequences on antiSMASH. Similarly, the BGCs for the structurally related rhodopetrobactin and methylolanthanin were obtained from the genomes of *R. palustris* TIE-1 (NCBI:txid395960) and *M. extorquens* AM1 (NCBI:txid272630). Additionally, genes in the petrobactin cluster were queried against *Microbulbifer* sp. CNSA002 using the tblastn functionality in NCBI’s BLAST^149^ (**Table S7**). BGC boundaries were determined using RAST-SEED predictions.

#### Direct metal infusion on UHPLC-MS/MS and data analysis

Relative binding assays were analyzed using a Thermo Scientific Q Exactive HF with upstream Vanquish UHPLC with a Kinetex 1.7 μm EVO C18 column (50 x 1.0 mm, 100 Å pore size) (Phenomenex; Torrance, CA, USA). A 5 uL injection was subjected to chromatographic separation at a flow rate of 0.150 mL/min with post-column metal infusion at 0.005 mL/min using a syringe pump. A 0.150 mL/min 10 mM ammonium acetate stream was fed using a PEEK T-splitter post-LC to keep the resulting stream at a pH ∼6^54^. Chromatographic separation was performed with the following mobile phases and gradient: A (H_2_O + 0.1% (v/v) formic acid) and B (acetonitrile + 0.1% (v/v) formic acid) with a linear gradient from 2.0 % B to 99.0% B over 6 min, held for 2 min as a washout, re-equilibrated by returning to 2.0 % B by 8.1 min, and held until 12 min. MS spectra were acquired in positive ionization mode from *m/z* 350-1500 Da for petrobactin and derivatives and 250-1500 Da for bulbichelin and derivatives (in order to observe the [M+2H]^2+^of Bulbichelin C) to 1500 Da (MS1 resolution of 45,000) and 100 ms maximum injection time with an AGC target of 1E6. The five most intense ions per MS^1^ were selected for fragmentation and MS^2^ data acquisition with a resolution of 15,000, and isolation window of 1.0 m/z. A stepped normalized collision energy of 25, 35, and 45 eV was used for fragmentation. Intensity threshold for selected ions was set to 1.6E5 and precursors were selected using an apex trigger of 2 to 15 s from first detection. The MS/MS dynamic exclusion parameter was set to 5 s. MS^2^ maximum injection time was 50 ms with an AGC target of 5E5 and minimum of 8E3. All data were acquired with the following parameters: sheath gas flow of 35 arbitrary units (AU), auxiliary gas flow of 10 AU, and sweep gas flow of 1 AU. Additionally, the auxiliary gas heater temperature was 200 °C, spray voltage was 3.5 kV, the capillary temperature was 250 °C, and the S-lens RF level was set to 55 AU. Metal solutions for infusion were prepared fresh on the day of use with LCMS-grade water and filtered prior to loading into the syringe. FeCl_3_ was used to provide Fe^3+^ ions, ZnCl_2_ for Zn^2+^ ions, CoCl_₂_·6H_₂_O for Co^2+^ ions, MnCl_2_·4H_2_O for Mn^2+^ ions, NiCl2·6H2O for Ni2+ ions, and (where applicable) CuSO_4_·5H_2_O for Cu^2+^ ions. All metal solutions were prepared at concentrations of 2 and (where applicable) 0.5 mM.

Raw data files were converted to the mzML format using msConvert by Proteowizard as previously described^54^. MS data was processed with mzmine 4.7.29 to produce feature tables from the data files. Peak-picking was performed with *m/z* 10 ppm tolerance and retention time with 0.16 min tolerance. A table of calculated and found *m/z* and retention time for each [M+H]^+^ and metal-adduct was prepared from the original feature table (**Tables S10-12**). A binding percentage, calculated as the metal-adduct peak area divided by the total peak area of the metal adduct plus the [M+H]^+^ adduct in each infusion run, was calculated for each infused metal and concentration (**Table S13**). Metal-adducts were not searched for if [M+H]^+^ species was not found. Bar graphs were prepared using all replicates (n = 3) for each infusion using the Plotly package for python with standard deviations shown as error bars.

## Supporting information

Siderophore structure examples, MS^2^ and NMR analysis of acylated petrobactins, siderophore moieties used for MassQL queries, and *Microbulbifer* sp. CNSA002 genomic analysis.

## Acknowledgements

This work was supported by the NIH R35GM150870 to N.G. and by NIH R35GM155026 to A.A. We thank Dr. Vinayak Agarwal Georgia Institute of Technology for sharing their bacterial strains with us.

## Author contributions

MM and NG designed the study and wrote the paper. MM did the experimental work and generated all figures. AA and CB conducted metal binding assays using metal infusion mass spectrometry (Figures 4C and 7C). All authors edited the manuscript. HW aided in NMR data acquisition.

## For Table of Contents Only

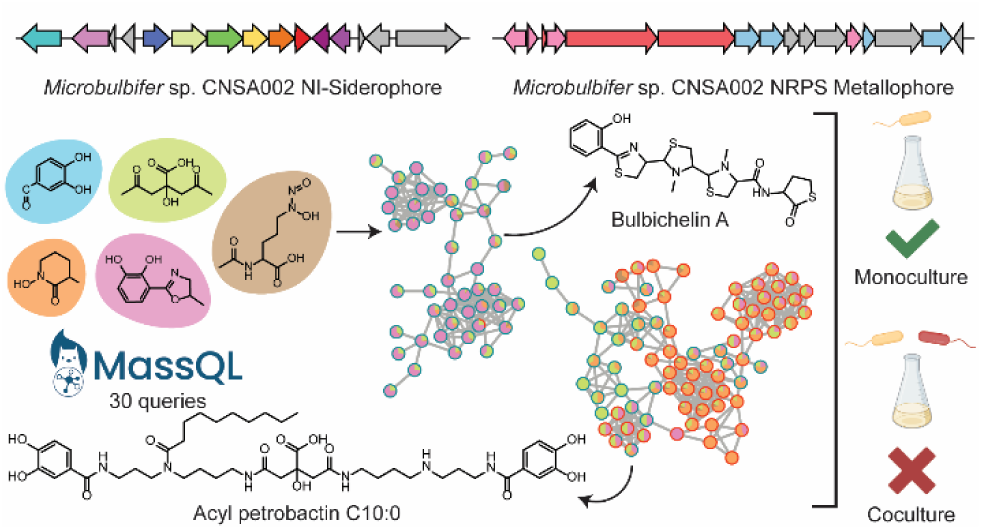

## Notes

### Competing Interest Statement

The authors have declared no competing interest.

